# Genome-wide DNA bridging by H-NS reshapes the stationary phase nucleoid and transcriptional landscape

**DOI:** 10.1101/2025.10.14.682352

**Authors:** Lindsey E. Way, Xiaofeng Dai, Emma E. Wiesler, Lauren A. McCarthy, Zhongqing Ren, Gail G. Hardy, David E.H. Fuller, Labeeb Hossain, Anne S. Meyer, Elio A. Abbondanzieri, Julie S. Biteen, Xindan Wang

## Abstract

Bacterial nucleoid-associated proteins (NAPs) structure the chromosome and regulate gene expression, but how these two functions are related is unclear. H-NS is a well-studied NAP that acts as a global gene silencer capable of bridging and looping DNA *in vitro*. Here, using high-throughput chromosome conformation capture assays, we show that H-NS mediates genome-wide long-range DNA looping in the stationary-phase nucleoid of *Escherichia coli*. Chromatin immunoprecipitation assays demonstrate that high levels of H-NS are present at the base of DNA loops. Super-resolution imaging and single-particle tracking show that H-NS binds more tightly in stationary phase and compacts the nucleoid mesh. Transcriptomic analyses indicate H-NS represses gene expression more strongly in the looped nucleoid and enables higher expression of genes outside of H-NS-bound regions. Overall, our study demonstrates that H-NS bridges distal DNA regions along the genome upon nutrient limitation, causing reduced nucleoid accessibility, stronger transcriptional repression, and a shifted transcriptional landscape.

## Introduction

The bacterial genome is dynamically structured by a complex network of proteins and cellular processes: nucleoid-associated proteins (NAPs) structure DNA and regulate gene expression; highly transcribed genes generate chromosomal interaction domains (CIDs); and DNA replication and transcription combined with topoisomerases change DNA supercoiling dynamics^1–5^. Together, these factors compact the genome by over 1000-fold, letting it fit within the cell while also allowing vital cellular processes to occur throughout the cell cycle.

DNA-bound NAPs form distinct architectures by bending DNA, wrapping DNA, bridging DNA loci, or stiffening DNA^1–5^. These activities restructure the genome and affect transcription. H-NS, a well-studied NAP found in γ-proteobacteria^6–9^, binds AT-rich DNA sequences where it acts as a global regulator to repress gene expression^6,7,9–12^. H-NS and H-NS-like proteins have been found to repress foreign or xenogeneic DNA in their respective bacteria, preventing RNA polymerase from transcribing these regions to generate toxic proteins that can negatively impact cell health and viability^6,10,13–17^. It has been well characterized that H-NS can form DNA bridges *in vitro* through radiolabeled DNA-bridging assays^6,18^, single-particle tracking experiments^18–20^, electrophoretic mobility shift assays^6,17^, and atomic force microscopy^17,19,21,22^. These studies have informed a mechanistic model for DNA bridging where H-NS dimers bind DNA and cooperatively multimerize to form a linear filament along DNA^19,23,24^, which then undergoes a conformational change that allows the multimer to bind a second segment of DNA, bridging the two DNA duplexes and forming a DNA loop^18,19^.

While H-NS bridging of DNA is well supported *in vitro*, *in vivo* evidence has been lacking. Genome-wide chromosome conformation capture techniques (3C-seq, Hi-C) have provided a map of short-range and long-range DNA interactions throughout the genome^25–28^. A recent ultra-high-resolution Micro-C analysis reported that *Escherichia coli* H-NS mediates the formation of chromosomal hairpins by bridging short regions of DNA, which have a median length of 2 kb^29^. In *Acinetobacter baumannii,* H-NS was found to cause a 3-kb DNA loop^6^. However, it has been unclear whether H-NS can mediate longer-range DNA bridging beyond 3 kb. A study in *Bacillus subtilis* observed enhanced DNA looping in stationary phase and found that the atypical H-NS-like protein Rok mediates these interactions^30–32^.

Here, we show that H-NS mediates genome-wide long-range DNA loops by bridging DNA *in vivo*. Using next-generation sequencing techniques, as well as super-resolution imaging and single-particle tracking, we analyze the characteristics and biological functions of H-NS-mediated DNA bridging in chromosome organization, nucleoid accessibility, and gene expression. We found that H-NS-mediated DNA loops reduce nucleoid accessibility, enable stronger transcriptional repression, and shift the transcriptional landscape globally.

## Results

### H-NS is required for bridging distant DNA loci in the *E. coli* nucleoid

Our recent work examined the folding patterns of the *E. coli* chromosome across growth phases using Hi-C experiments^33^. During deep stationary phase, occurring after 96 h of growth in batch culture, we found that the *E. coli* chromosome is restructured and shifts from a characteristic plaid pattern^29^ to a checkerboard pattern of gridded “dots”^33^ (Figs. 1A-B, black arrows). This checkerboard pattern indicates that distal DNA loci are interacting with each other and generating long-range, point-to-point DNA contacts in the form of DNA loops (Fig. 1G, left). To test whether H-NS mediated these interactions, we performed additional Hi-C experiments in *Δhns.* We found that chromosomal looping was abolished in deep stationary phase, indicating H-NS is required for DNA looping (Fig. 1C). We next sought to determine whether other NAPs play a role in chromosomal looping. Previous studies showed that the *E. coli* NAP StpA can form heterodimers with H-NS, and StpA binds DNA with a profile very similar to H-NS^34^. Additionally, deletion of H-NS has been shown to increase StpA expression levels^35^. Interestingly, the Hi-C map of *ΔstpA* at 96 h demonstrated that DNA looping was still present and appeared even stronger than in the wild-type (WT) (Fig. 1D), showing that StpA is not required for chromosomal looping. In addition, the deletion of other NAPs (*Δfis* and *ΔihfAB)* showed enhanced DNA looping at 96 h (Figs. 1E-F). These results show that H-NS is required for bridging distal DNA loci and restructuring the *E. coli* nucleoid in deep stationary phase, while the presence of other NAPs reduces H-NS-mediated DNA looping.

**Figure 1.**
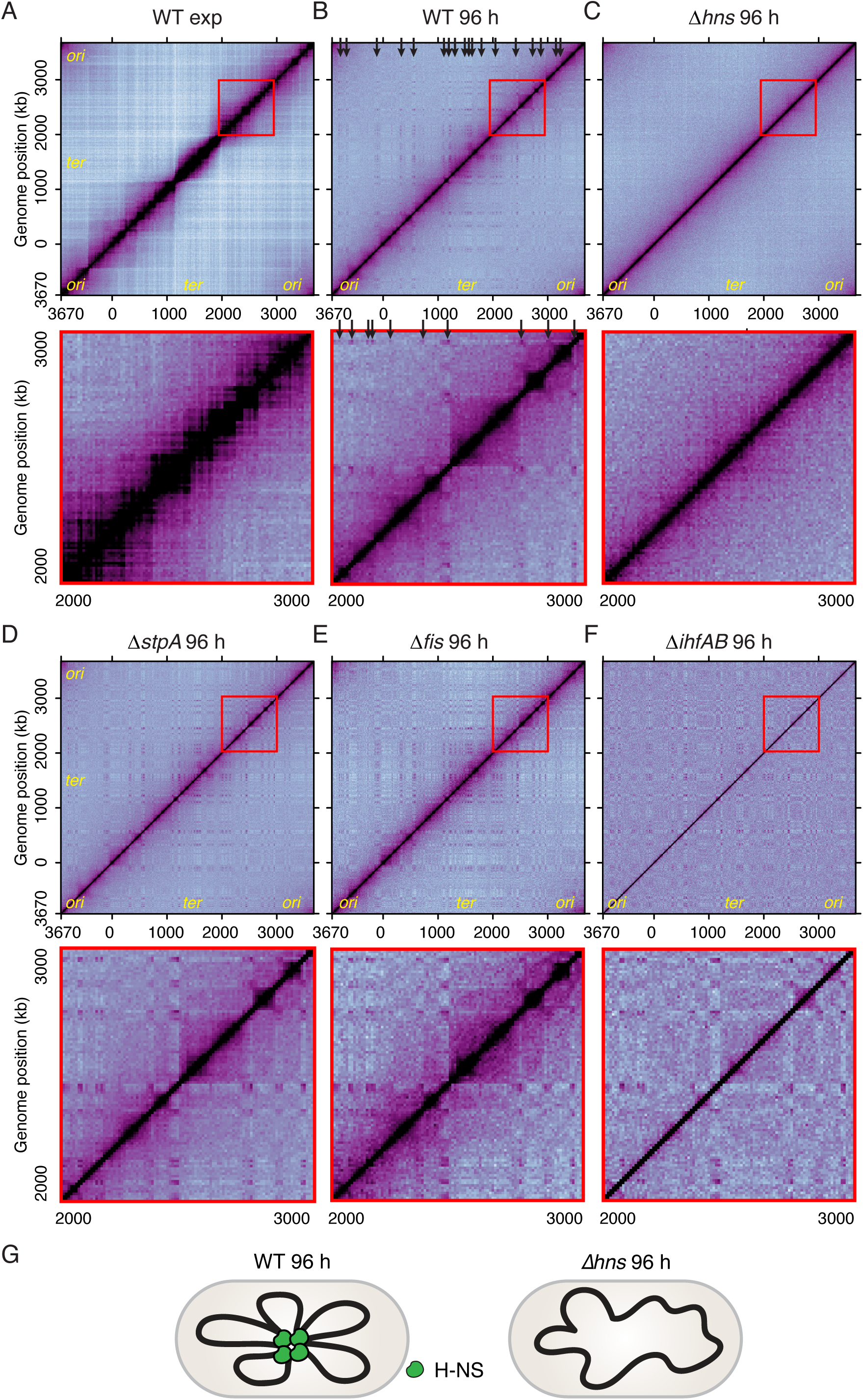
H-NS is required for DNA looping in deep stationary phase. **(A-F)** Hi-C maps of indicated strains and conditions: WT grown in exponential phase (A) and deep stationary phase (96 h) (B); *Δhns* (cWX2870) (C); *ΔstpA* (cWX3338) (D); *Δfis* (cWX2478) (E); or *ΔihfAB* (cWX2943) (F) grown to deep stationary phase. Top panels show whole genome maps using a bin size of 10 kb. The 1000-kb area in the red box is zoomed in on the bottom panels. Black arrows point to examples of gridded dots. **(G)** Schematic model.

### H-NS is highly enriched at DNA-loop anchors

To quantitatively analyze the chromosome folding patterns in exponential phase and stationary phase, we developed a fully automated image processing algorithm to isolate the sharp dots associated with the bridged loci that anchor chromosomal loops. Briefly, we used a rolling ball background subtraction method^36^ to remove broad features (>100 kb) of the Hi-C plots, then applied an automated threshold to remove the intense central diagonal signal from the plots. The processed Hi-C plot was integrated in one dimension to identify Hi-C peaks, which represent regions of the chromosome that come together to anchor the DNA loops (see Method Details) (Fig. S1). Applying this algorithm, strong peaks representing DNA-loop anchors emerged in deep stationary phase (Figs. 2A-B) and these peaks were completely abolished by deletion of *hns* (Fig. 2C). We identified 28 regions on the Hi-C map that strongly interact with each other (Fig. 2B bottom panel, Supplementary Dataset 1). These interactions were not detected in the *Δhns* strain (Fig. 2C) but were enhanced in the absence of StpA, Fis, or IHF (Fig. S2).

**Figure 2.**
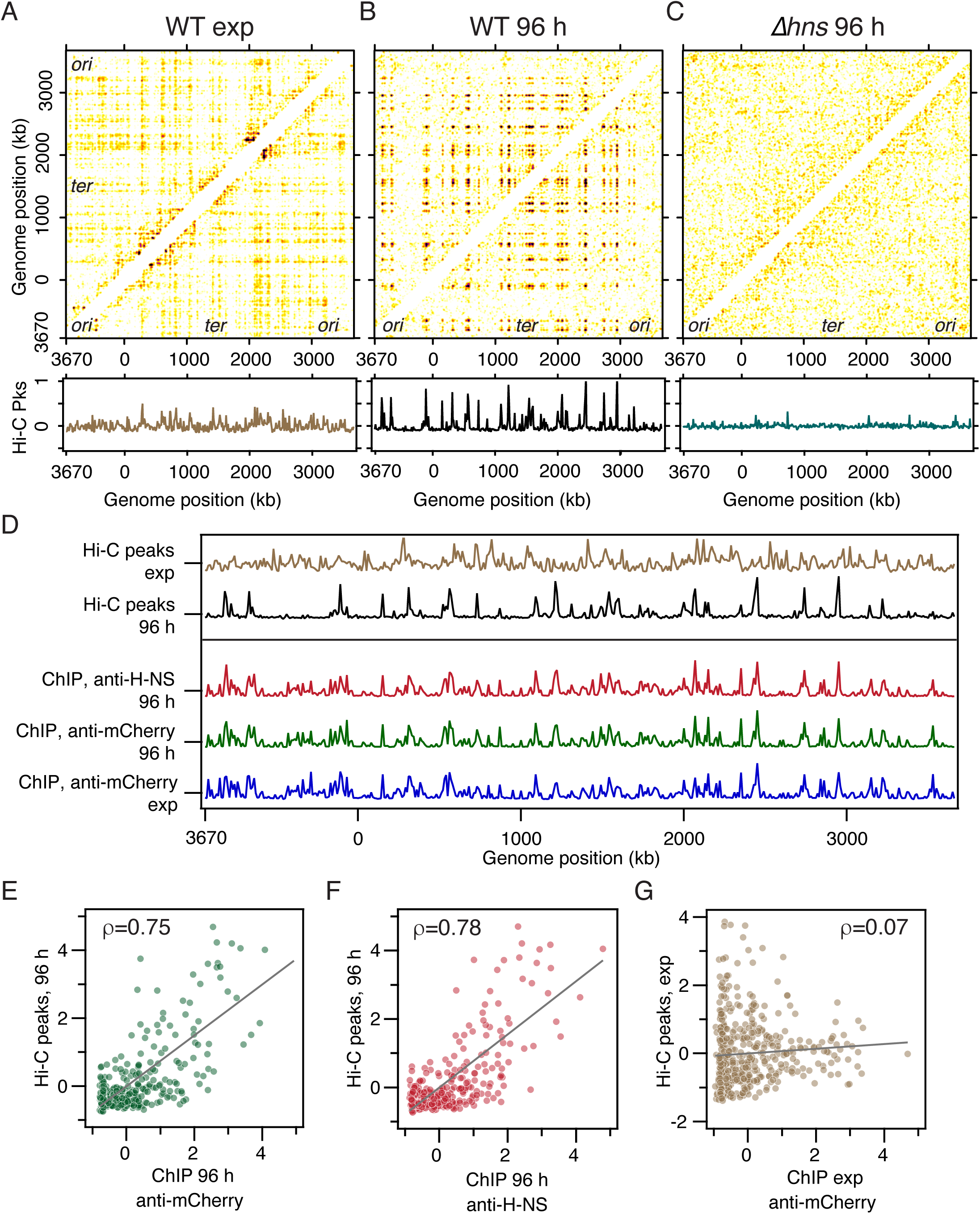
Hi-C loop anchors are correlated with high H-NS ChIP-seq enrichment. **(A-C)** Analysis of Hi-C loop anchors using a rolling ball background subtraction algorithm for WT cells at exponential phase (A) and deep stationary phase (B), and *Δhns* (cWX2870) at deep stationary phase (C). Top panels show the genome-wide, pairwise distribution of narrow peaks representing regions that are forming DNA-loop anchors. Bottom panels show the linear integrated density of the peaks across the genome. See also Method Details and Fig. S1. **(D)** Comparison of Hi-C peaks and ChIP-seq peaks. Hi-C peaks in (A-B) were normalized and replotted to compare with 10-kb-binned ChIP-seq results using anti-H-NS antibodies in WT (deep stationary phase) or anti-mCherry antibodies in cells expressing H-NS-PAmCherry (cWX2882) (exponential phase and deep stationary phase). **(E-G)** Scatter plots showing the Pearson correlation of Hi-C peaks and ChIP-seq peaks between different growth conditions. Strains used are cWX2882, cWX2200, and cWX2882.

We then used chromatin immunoprecipitation sequencing (ChIP-seq) to characterize the genome-wide distribution of H-NS. We performed ChIP-seq in WT and *Δhns* growing at exponential and deep stationary phases using anti-H-NS antibodies^37^ (Figs. S3A-B). Separately, since anti-H-NS antibodies are known to be unspecific^37^, we used a strain expressing *hns-PAmcherry*^38,39^ at the endogenous *hns* locus and performed anti-mCherry^40^ ChIP-seq at exponential and deep stationary phases (Figs. S3C-D) to validate the ChIP-seq results. We found that at deep stationary phase, anti-H-NS antibodies against untagged H-NS and anti-mCherry antibodies against H-NS-PAmCherry have overlapping ChIP-seq profiles (Fig. S3E), indicating that both results are valid. At exponential phase, only H-NS-PAmCherry ChIP-seq results are usable due to antibody specificity (Figs. S3A-D, right panels). We then compared anti-mCherry ChIP-seq results between exponential and deep stationary phases and found that H-NS has essentially overlapping enrichment profiles at these two growth phases (Fig. S3F).

To understand whether H-NS is enriched at the DNA-loop anchors, we replotted the H-NS ChIP-seq data using 10-kb bins and compared the results to the Hi-C peaks identified above (Fig. 2D). All datasets were normalized to have a mean of zero and standard deviation of 1 to aid the comparison. For the 96-h time point (deep stationary phase), we used ChIP-seq results from both anti-H-NS and anti-mCherry antibodies; for exponential phase results, we used only anti-mCherry ChIP-seq data. The plots showed that Hi-C peaks from 96 h aligned with peaks in the ChIP-seq data (Fig. 2D). Correlation analysis revealed that during deep stationary phase, ChIP-seq and Hi-C peaks were strongly correlated (ρ ≥ 0.75, Figs. 2E-F). At exponential phase, ChIP-seq and Hi-C peaks were weakly correlated (ρ = 0.07, Fig. 2G). These results indicate that H-NS directly mediates DNA looping during deep stationary phase but not during exponential phase.

Interestingly, not all H-NS ChIP-seq peaks overlapped with Hi-C-identified DNA-loop anchors (Fig. S4A). When focusing only on bins that were more than one standard deviation above the mean, a large majority of deep stationary phase Hi-C peaks (83%) overlapped with the H-NS peaks, but just over half (58%) of H-NS peaks overlapped with the Hi-C peaks (Fig. S4B). To understand this discrepancy, we analyzed characteristics of H-NS ChIP-seq peaks that overlapped with the DNA-loop anchors. In each Hi-C peak, we found a cluster of H-NS enrichment peaks spanning a long region (Supplementary Dataset 1). At a higher resolution (1-kb), these clusters had an average of 3.6 distinct peaks that spanned an average region of 35.8 kb (see two examples in Fig. S3E middle and right panels). The distinct peaks within each cluster had variable width; the widest single peak of each cluster spanned an average region of 11.3 kb (Supplementary Dataset 1). In comparison, for regions containing H-NS peaks that did not overlap DNA-loop anchors (see an example in Fig. S3E left panel), H-NS ChIP-seq peaks had an average of 2.0 peaks per cluster and spanned an average of 12 kb, with the widest single peak spanning 4.6 kb (Supplementary Dataset 1). Collectively, our results show that longer stretches of DNA with higher levels of H-NS occupancy correlate with DNA-loop anchors, suggesting that a high density of H-NS is required to bridge DNA and form DNA loops.

### H-NS molecules diffuse more slowly in deep stationary phase

Our Hi-C analysis indicated that H-NS mediates DNA looping in deep stationary phase but not in exponential phase (Fig. 1A-B). Our ChIP-seq results showed that H-NS enrichment peaks were nearly identical at both timepoints (Figs. 2D, S3F), leading us to conclude that H-NS distribution is not the only factor driving DNA-loop formation. To understand whether H-NS binds more tightly to DNA during deep stationary phase to allow DNA-loop formation, we quantified H-NS dynamics in live cells. We tracked single H-NS-PAmCherry molecules using photoactivated localization microscopy (PALM) super-resolution imaging^41^ (Fig. 3) in the same strain used for ChIP-seq experiments (Fig. S3). Although the PAmCherry tag has been shown to induce aggregation under certain conditions^39^, our ChIP-seq experiments showed that H-NS-PAmCherry and untagged H-NS had the same binding profile at deep stationary phase (Fig. S3E). Importantly, our Hi-C results showed that although the chromosomal looping patterns were weakened, there were still detectable DNA loops in the H-NS-PAmCherry strain (Fig. S3G), indicating that H-NS-PAmCherry has retained the function of endogenous, untagged H-NS.

**Figure 3.**
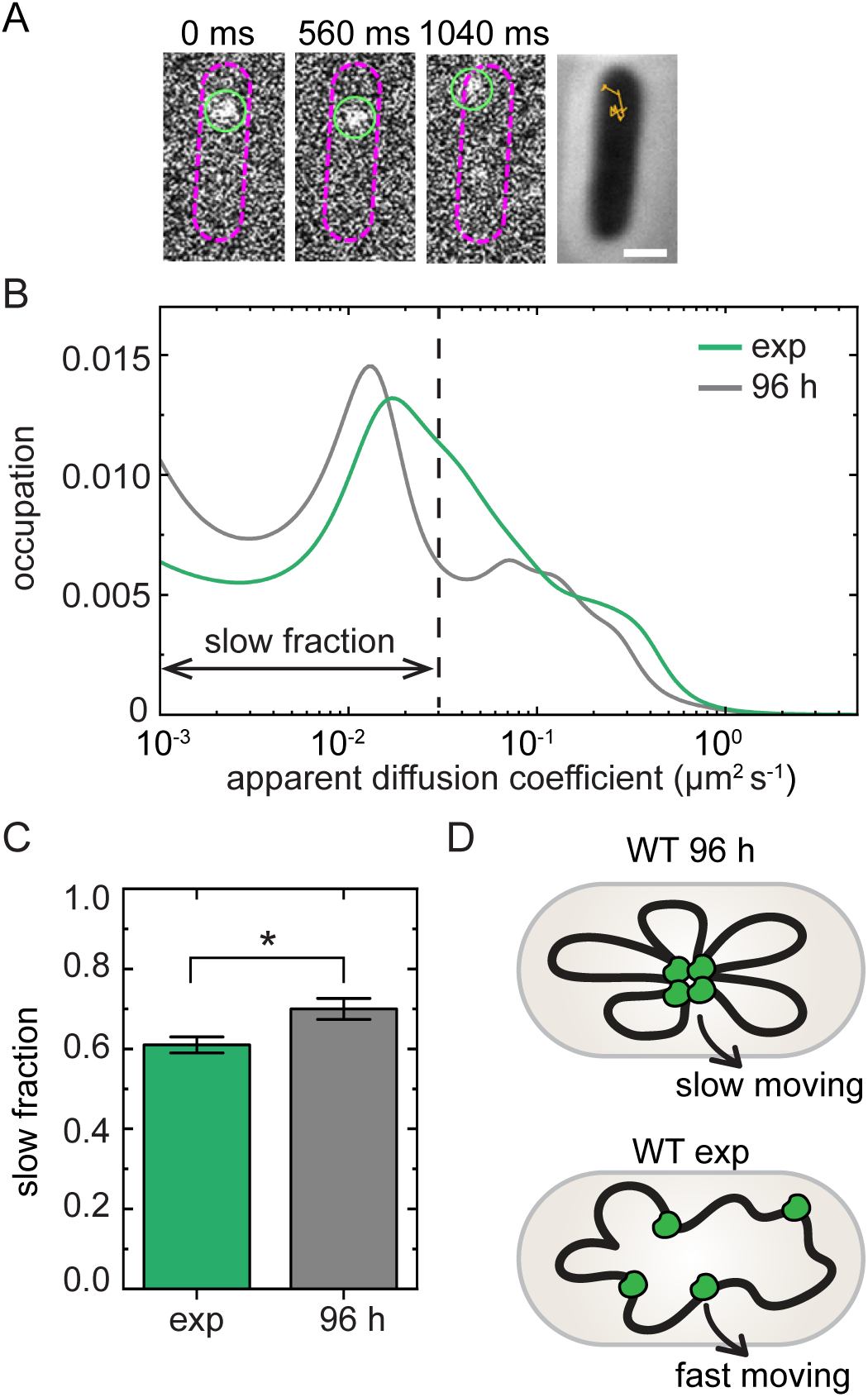
Dynamics of single H-NS-PAmCherry molecules in the exponential and stationary phases. **(A)** Representative fluorescence images of a single H-NS-PAmCherry molecule that is localized (green circles) in each frame acquired every 40 ms and tracked from frame to frame (cWX2882). Smoothed cell outlines are shown in pink. Phase-contrast image of an exponential-phase cell overlaid with the trajectory of motion for the same H-NS-PAmCherry molecule is shown in the right panel. Scale bar, 1 µm. **(B)** Apparent diffusion coefficient distributions of H-NS-PAmCherry molecules measured for three biological replicates for each growth phase and analyzed using the State Array Bayesian framework^66^. The dashed vertical line represents the limit of resolution for slowly diffusing molecules considering our localization precision and exposure time; molecules with diffusion coefficients below this cutoff were assigned as “slow”. Three biological replicates were analyzed. Plots for each individual replicate can be found in Fig. S5. **(C)** Average fraction of slow molecules for the exponential and stationary phases. Error bars are the standard deviation from three biological replicates. The star indicates that the mean slow fractions between the two growth phases are significantly different (*p* < 0.05) by a two-sample t-test. **(D)** Schematic model.

Single-molecule tracking experiments were performed by photoactivating ∼1-4 H-NS-PAmCherry molecules per cell at a time, localizing those molecules in each frame of a fluorescence movie, and compiling trajectories of motion (Fig. 3A). We repeated this activation and tracking process dozens of times per cell for ≥ 20 cells per condition, collecting ∼2000 trajectories for each growth phase. Analysis of the H-NS-PAmCherry trajectories showed that, during deep stationary phase, H-NS molecules had a slower peak diffusion coefficient than in exponential phase (Fig. 3B). We determined the slow-moving fraction of molecules for each growth phase by assigning a slow fraction cutoff based on our localization precision and exposure time. We found that on average, deep stationary-phase H-NS molecules were significantly more likely to diffuse slowly compared to exponential-phase H-NS molecules (Fig. 3C). This significant difference in H-NS dynamics between the exponential and stationary phases was observed across three biological replicates (Fig. S5). These results are consistent with our model that H-NS binds tightly and forms stable DNA bridges between distal loci in deep stationary phase, slowing the diffusion of H-NS relative to exponential phase (Fig. 3D).

### H-NS-mediated chromosome looping reshapes the transcriptional landscape

Next, we examined whether chromosome looping affects transcriptional regulation. We performed RNA-sequencing in WT and *Δhns* strains growing in exponential and deep stationary phases. Three biological replicates for each sample were analyzed and showed highly reproducible results (Fig. S6A). We calculated the differential expression of each gene (i.e., expression fold change, or FC) in WT vs. *Δhns* (Figs. S6B-C), then used the EdgeR package to calculate raw p-values and adjust them for false discovery rate (FDR)^42^. Volcano plots of FDR vs. fold change (FC) showed that many genes had significantly altered expression by H-NS (Fig. 4A-B blue and red dots). While a similar number of genes were significantly downregulated by H-NS (FDR < 0.05 and FC < 0.5) in both exponential phase and at 96 h (840 vs. 689, Figs. 4A-B blue dots), fewer genes were significantly upregulated (FDR < 0.05 and FC > 2) in exponential phase compared to deep stationary phase (117 vs. 559, Figs. 4A-B red dots).

**Figure 4.**
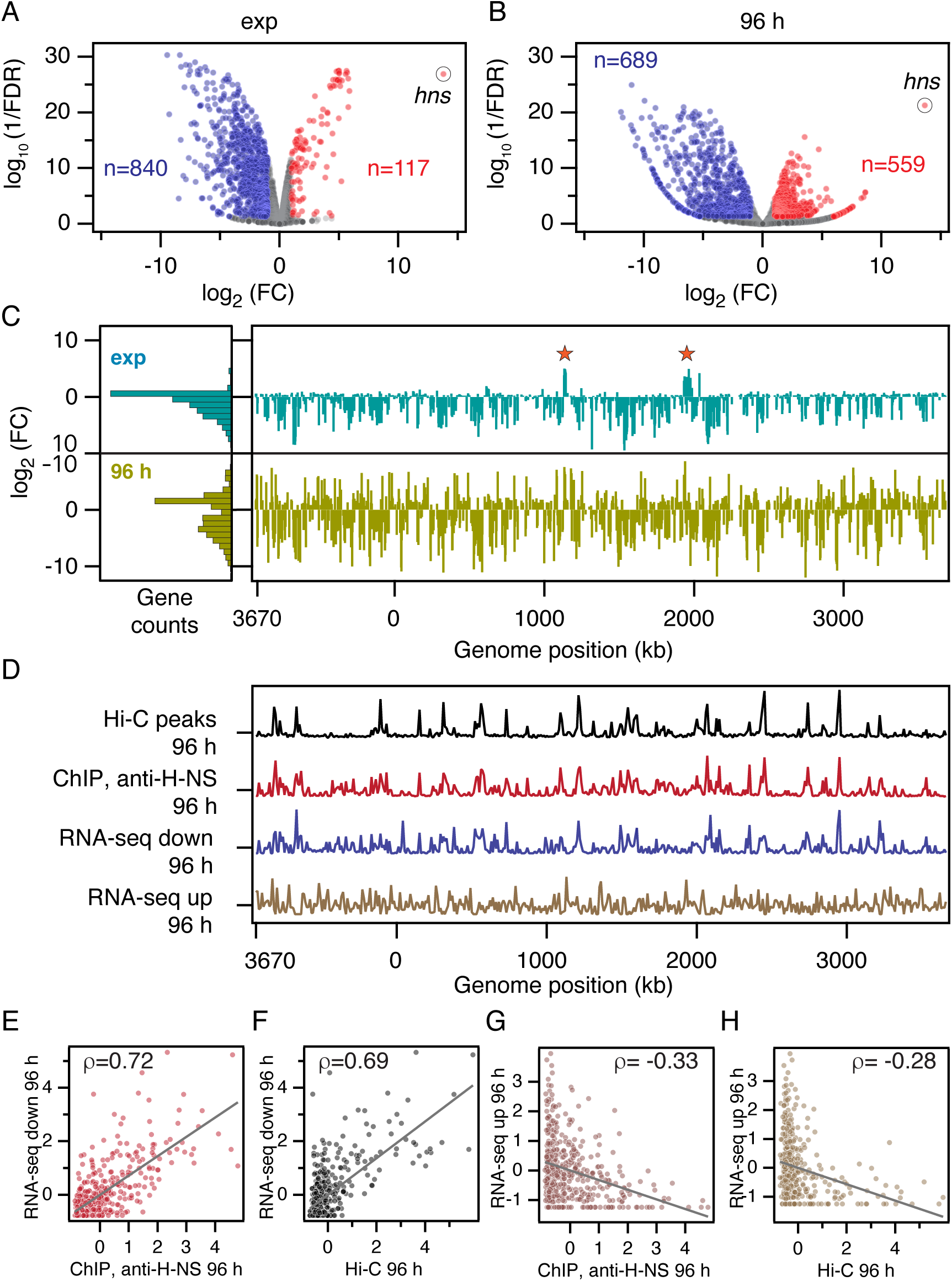
Transcription repression is correlated with Hi-C loop anchors and H-NS ChIP-seq enrichment. **(A-B)** Volcano plots showing differential expression of genes in WT and *Δhns* (cWX2882) at exponential phase (A) and deep stationary phase (B). For each gene, the mean expression in the WT strain was divided by the corresponding value in the *Δhns* strain to calculate fold change (FC = WT*/Δhns*). The *x*-axis represents the log2 of FC. The false discovery rate (FDR) represents the threshold required to consider an observed change to be significant when taking into account multiple hypothesis testing. The *y*-axis represents the 1/FDR in log10 scale. Gray dots represent FDR > 0.05 or |log2(FC)|< 1. For genes with an FDR < 0.05, blue and red dots represent log2(FC) < −1 or >1, respectively. The *hns* gene is highlighted. Number (*n*) of downregulated (blue) or upregulated (red) genes are shown. (C) Right panel: log2(FC) for genes that had *p* < 0.05 at exponential phase and deep stationary phase. The x-axis shows the position of each gene’s transcription start site. Left panel: aggregated values in histograms. Orange stars indicate genes that are upregulated by H-NS at exponential phase. (D) Comparison of Hi-C, ChIP-seq, and RNA-seq results in 10-kb bins at deep stationary phase. For the RNA-seq results, a weighted histogram of downregulated and upregulated genes is shown (see Method Details). **(E-H)** Scatter plots showing the Pearson correlation of the RNA-seq histograms against the ChIP-seq peaks and Hi-C peaks. Strains used are cWX2200 and cWX2870.

To examine the genetic locations of upregulated and downregulated genes, we plotted FC against transcription start sites (TSS) for every gene with a *p*-value < 0.05 for both the exponential and deep stationary phase datasets (Fig. 4C). We found that in exponential phase, highly upregulated genes were mostly limited to two narrow clusters (Fig. 4C, top panel, orange stars) and were mainly components of the flagellum and flagellar motor (Supplementary Dataset 2). This result is consistent with previous findings that motility genes are upregulated by H-NS through the downregulation of *csgD*^43^, which we also observed (FC = 0.08, FDR = 1.6 × 10^-16^). In contrast, during deep stationary phase, highly upregulated genes were widely distributed across the genome (Fig. 4C, bottom panel).

To compare the pattern of differential gene expression across the genome with our Hi-C and ChIP-seq results, we created a weighted histogram of all downregulated genes in deep stationary phase (Fig. 4D, blue trace). Within each 10-kb bin, we summed the negative log_2_ of FC for each gene with an FC below 0.5 to estimate the strength of downregulation in that region. The resulting histogram was normalized and compared to the normalized plots of our Hi-C and ChIP-seq peaks (Fig. 4D). We found that the downregulated genes were strongly correlated to H-NS binding (ρ = 0.72, Fig. 4E) and DNA-loop anchor loci (ρ = 0.69, Fig. 4F). A more moderate correlation was observed between downregulated genes and H-NS binding or Hi-C peaks (ρ = 0.58 and 0.20, respectively) in exponential phase (Fig. S6D), when strong DNA looping was not observed. Overall, during deep stationary phase, DNA-loop anchor loci were correlated with both H-NS enrichment and repressed gene expression (Fig. S6E).

For both exponential and deep stationary phase samples, we found that the locations of downregulated genes were correlated with H-NS-enriched loci, but the correlation was stronger and the fold change was larger during deep stationary phase (Figs. 4C-D, S6D), indicating that H-NS represses these genes more effectively when it bridges these DNA. In addition, we found that more genes were significantly upregulated by H-NS in deep stationary phase than in exponential phase (559 vs. 117, Figs. 4A-B). These genes were distributed throughout the genome (Fig. 4C bottom panel) and were concentrated outside of H-NS-bound regions. A weighted histogram of upregulated genes during deep stationary phase, generated by summing log_2_(FC) for every gene with an FC > 2, was weakly anti-correlated to H-NS binding and DNA-loop anchor loci (ρ = −0.33 and −0.28, respectively, Figs. 4G-H). We conclude that in deep stationary phase, H-NS strongly represses gene expression in the regions it binds, and enhances gene expression in regions it does not bind.

### DNA bridging tightens the nucleoid mesh

Our Hi-C, ChIP-seq, and RNA-seq results are consistent with a model in which high levels of H-NS bridge DNA to form a rosette structure in stationary phase (Fig. 1G) and strongly repress gene expression in bound regions. We examined whether H-NS affects the physical structure of the nucleoid and modulates its accessibility to large protein complexes. To address this question, we used single-particle microscopy to track 25-nm eGFP-labeled protein nanocages in living *E. coli* cells^44^ (Fig. 5A) and quantified their exclusion from the nucleoid (Fig. 5B). Because of their 25-nm diameter, these nanocages are a good analog of biomacromolecules like ribosomes^44,45^. These extrinsic tracers specifically measure the structure of the bacterial chromosome because they have no specific interactions with cytoplasmic cell components^46^.

**Figure 5.**
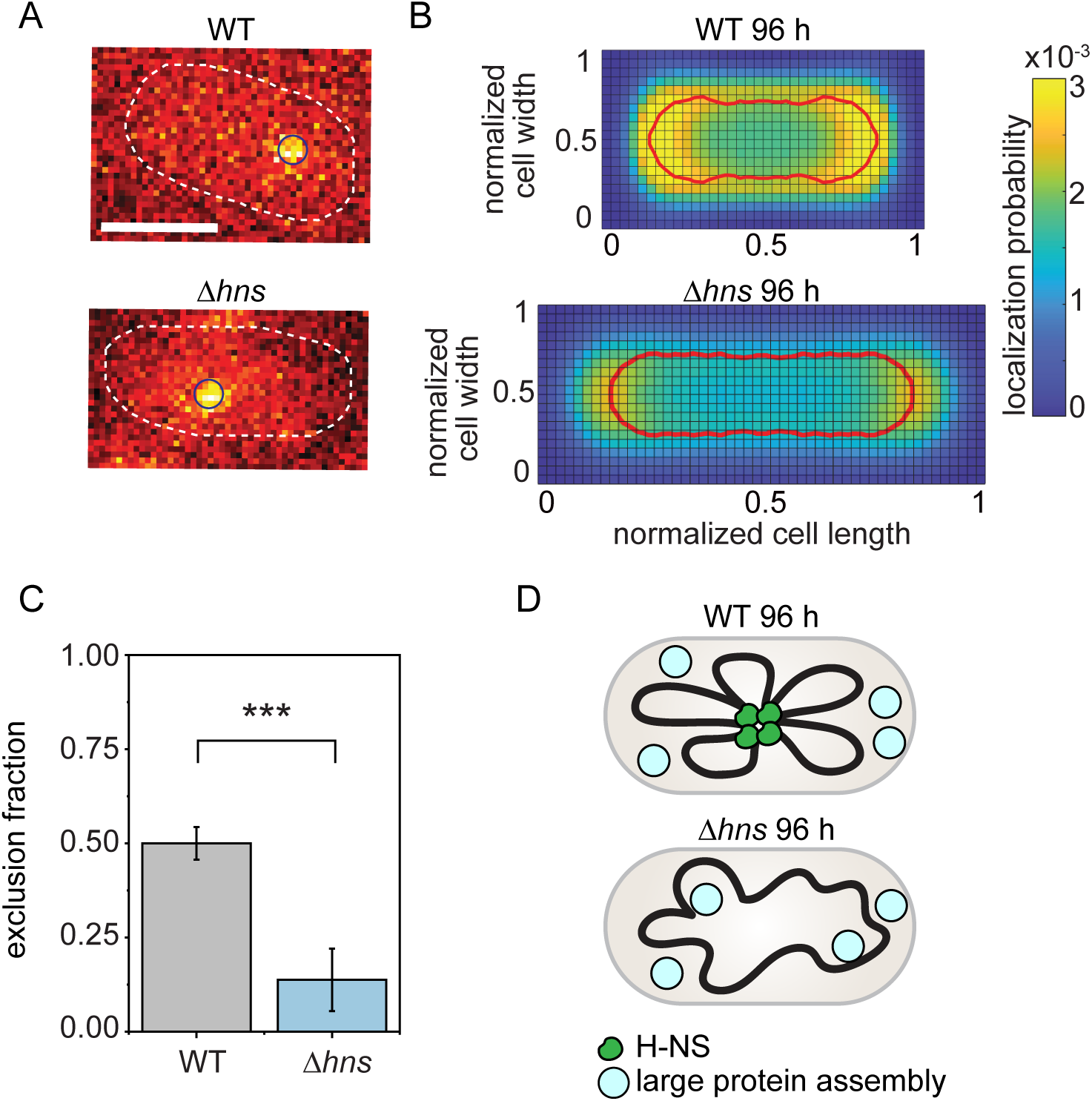
Exclusion of nanocages from the nucleoid in WT and Δ*hns* cells at deep stationary phase. **(A)** Representative fluorescence images of a single fluorescent nanocage that is localized (black circles) in each frame acquired every 40 ms (LAM044 and LAM065). Cell outlines from segmentation results are shown in white. Scale bar, 1 µm. **(B)** Nanocage localization heatmap of WT (LAM044) and Δ*hns* (LAM065) during deep stationary phase. Red lines: nucleoid outline from HUα-PAmCherry PALM imaging (Fig. S7A). **(C)** Normalized exclusion fraction of nanocages from the nucleoid. Bar heights represent the mean value of the normalized exclusion fraction from five random subsamples from *n* > 1500 cells within two biological replicates. Error bars show the standard deviation of the same process. Three stars represents a statistical difference (*p* = 0.0002) from a two-sample *t-*test. **(D)** Schematic model.

We tracked the nanocages in WT and Δ*hns* cells and combined the data from 7500 cells from each condition into a two-dimensional (2D) histogram (see Method Details) to map the average localization probability of the nanocages within the cells during deep stationary phase (Fig. 5B). To understand the position of nanocages relative to the nucleoid, we determined the nucleoid region in WT and Δ*hns* cells by PALM super-resolution imaging of HUα-PAmCherry^39^, which nonspecifically binds the chromosome^33,47,48^. For each condition, data from 30 cells were combined into a 2D histogram to determine the nucleoid boundary (Fig. S7A, red lines), which was overlaid on the nanocage result (Fig. 5B, red lines). We found that the nucleoid excluded nanocages in both WT and Δ*hns* cells, as indicated by the decrease in nanocage localization probability in the nucleoid region relative to the cytoplasmic region (Fig. 5B).

We then quantified the degree of nucleoid exclusion by accounting for the 3D cell shape (see Method Details) while considering the increased cell length of Δ*hns* cells (Figs. S7A-B). We normalized the exclusion fraction based on the computed theoretical minimum and maximum exclusion fractions for the average nucleoid and cell geometries in each condition. On this scale, an exclusion fraction of 1 represents complete exclusion by the nucleoid, meaning that all nanocages are located outside of the nucleoid in 3D, and an exclusion fraction of 0 indicates no exclusion by the nucleoid, meaning equal density of nanocages inside and outside the nucleoid in 3D. In deep stationary phase, we found increased exclusion of nanocages from the nucleoid in the WT compared to the *Δhns* mutant (Fig. 5C). This indicates that the absence of H-NS loosens chromosome structure, enhancing nucleoid accessibility (Fig. 5D), and supports a model where H-NS-mediated DNA looping tightens the nucleoid, restricting nucleoid accessibility and reshaping the transcriptional landscape in deep stationary phase.

## Discussion

H-NS has been studied for over three decades for its role in nucleoid structuring and gene silencing^4,6–9,13–15,18,20–22,37,49^. There has long been concrete evidence for H-NS performing DNA-bridging *in vitro*^17–22^. Very recently, *in vivo* evidence for H-NS-mediated DNA-bridging has been shown, but only for short ranges of 2 – 3 kb^6,29^. No reproducible *in vivo* evidence has been provided for long-range DNA-bridging mediated by H-NS. We have now shown that, during deep stationary phase, H-NS mediates genome-wide long-range DNA interactions by bridging dozens of loci *in vivo*.

Our ChIP-seq results demonstrated that H-NS has essentially the same binding profile during exponential and deep stationary phase. However, strong DNA bridging and looping are only observed during deep stationary phase, possibly for several reasons. First, active replication and transcription during exponential phase could disrupt long-range contacts in the genome, preventing H-NS from forming DNA loops. Secondly, topoisomerases and DNA translocases might cause dynamics in DNA that prevent stable DNA bridging during exponential phase. Finally, various NAPs, which have growth-phase-dependent expression levels^50^, could alter DNA interactions and prevent H-NS from forming DNA loops. Indeed, our analyses show that the absence of StpA, Fis, or IHF enhanced H-NS-mediated DNA bridging during deep stationary phase (Figs.1D-F and S2), consistent with the idea that other NAPs may prevent H-NS from forming stable DNA loops.

Our study offers particular insights into the interplay of StpA and H-NS *in vivo*. StpA is a paralog of H-NS that binds to similar DNA sequences and may share some overlapping functions^34,35,37^. In addition, StpA has been shown to form strong bridging interactions between different DNA molecules *in vitro* and can even enhance bridging by H-NS when both proteins are combined^51^, suggesting that StpA may act cooperatively with H-NS. However, a recent *in vivo* study that identified the formation of short-range chromosomal hairpins mediated by H-NS did not observe any relative enhancement of the structures by StpA^29^. Instead, the authors observed no change in hairpin formation in *ΔstpA*, weakened hairpins in *Δhns,* and no hairpin structures in *ΔhnsΔstpA*, suggesting that StpA is less effective at forming bridging contacts *in vivo*. In our study, we find no detectable long-range bridging interactions mediated by StpA when H-NS is deleted (Fig. 2C) and enhanced bridging interactions mediated by H-NS when StpA is deleted (Fig. S2), suggesting that StpA acts somewhat antagonistically to H-NS on these larger length scales. Taken together, these data point to a complex relationship between StpA and H-NS. We hypothesize that StpA forms weaker bridging interactions than H-NS *in vivo*, allowing it to partially substitute for H-NS for short-range interactions but not for the genome-wide DNA bridging reported here.

Although we do not know the causal relationship, we show that DNA bridging is correlated with slower H-NS movement, a more condensed nucleoid meshwork, lower DNA accessibility to protein complexes, stronger repression of H-NS-bound regions, and enhanced transcription of H-NS-free regions. Our results are consistent with a model where, during deep stationary phase, H-NS binds more tightly to the DNA, forming stable DNA bridges and tightening the nucleoid mesh, allowing H-NS-free regions to be better transcribed (Fig. 6).

**Figure 6.**
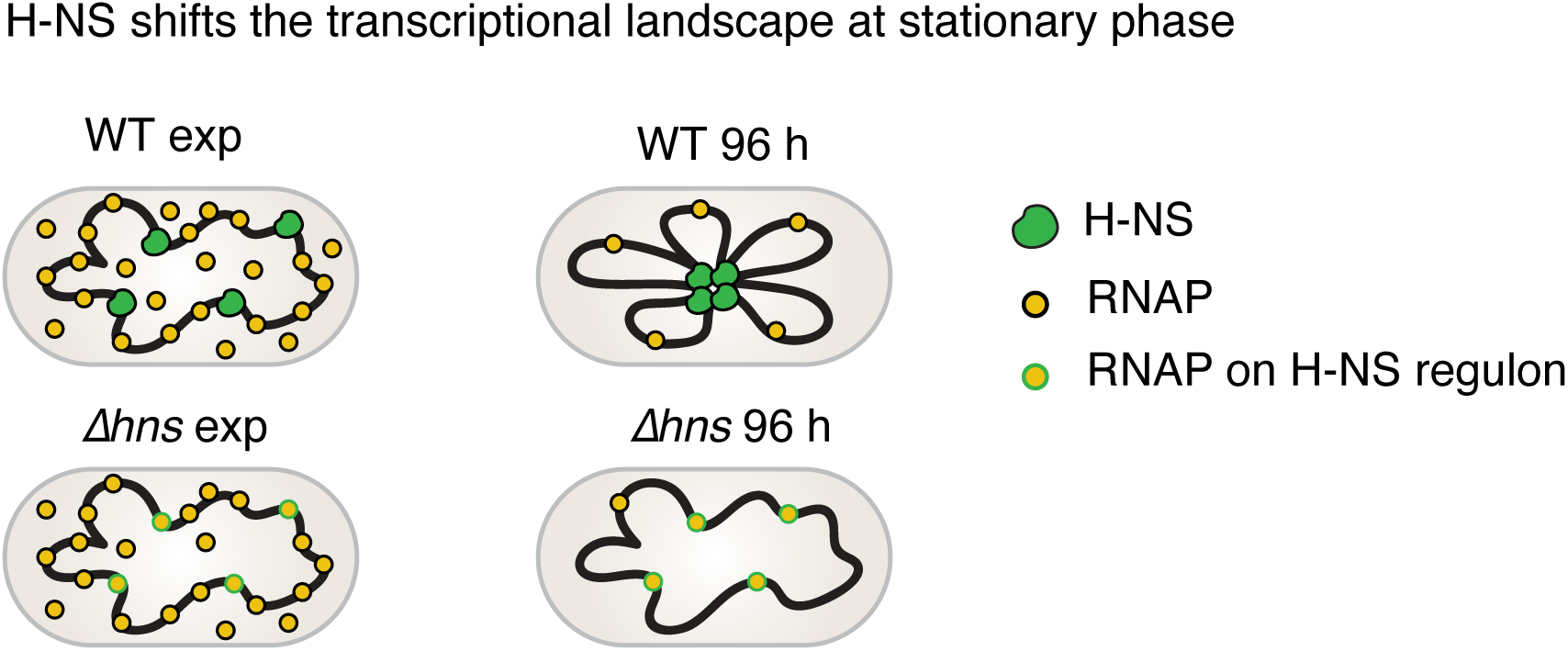
Schematic model. Left panel: at exponential phase, in the WT, H-NS binds to specific regions and represses the expression of the H-NS regulon; RNA polymerases (RNAPs) transcribe H-NS-free regions. In *Δhns*, RNAPs can express the H-NS regulon, but this does not affect H-NS-free regions because of the excess of RNAPs. Right panel: at 96 h deep stationary phase, in the WT, H-NS bridges specific regions of the nucleoid, tightening the nucleoid mesh and reducing RNAP’s access to the center of the nucleoid. This activity strongly represses the expression of the H-NS regulon and enables RNAPs to transcribe H-NS-free regions. In *Δhns*, RNAPs can access the H-NS regulon, which competes for the limited numbers of RNAPs and lowers the expression of H-NS-free regions.

H-NS-mediated DNA looping may be an effective strategy for cells to reshape the transcriptional landscape. During deep stationary phase, when nutrients are depleted, most RNA polymerase is sequestered in an inactive state^52^. Therefore, it becomes vital that all spurious transcription and translation of xenogeneic genes are tightly repressed to conserve limited resources^14,16^. We note that during exponential phase, H-NS primarily represses genes only where it binds, while in deep stationary phase, H-NS represses genes more strongly where it binds while also enhancing gene expression in regions it does not bind (Fig. 4C). This additional function of H-NS in stationary phase may be due to genome restructuring because genes buried near the center of the nucleoid will be less accessible to transcription and translation machinery while regions of DNA at the periphery of the nucleoid will be more accessible. The tighter mesh of the nucleoid caused by H-NS (Figs. 5B-C) should make this effect even stronger. The stronger repression of H-NS-bound genes may also be a direct result of H-NS bridging, since bridged H-NS filaments have been observed to pause RNAP and enhance transcription termination *in vitro*^17^.

A previous study in *B. subtilis* also observed DNA looping during stationary phase and found that it is mediated by the atypical H-NS-like protein Rok^30,31^. Thus, stable bridging of xenogenic sequences and physically sequestering them might be an important, conserved strategy for diverse organisms to efficiently allocate the limited supply of transcriptional and translational resources during stationary phase.

In summary, our study demonstrates that the master transcriptional regulator H-NS bridges distal DNA loci *in vivo* and reveals that a biological function of DNA bridging is the enhancement of gene silencing. We propose that this genome organization strategy optimizes survival during starvation by controlling the accessibility of different DNA regions to cellular machines like RNA polymerases. Our study highlights the importance of examining microbe behavior across different growth phases and stress conditions, since exponential phase measurements may not accurately reflect the entirety of roles played by a specific protein in the cell.

## Method Details

### *E. coli* strains and growth

Bacterial strains were derived from prototrophic *E. coli* K-12 strain W3110 (CGSC no. 4474)^53^. The growth medium we used was High-Def Azure medium (Teknova 3H5000) supplemented with 0.2% glucose (m/v), which we named HDA medium for short. Cells were grown at 37°C with shaking. For exponential phase timepoints, cells were grown until an OD_600_ of 0.2-0.5 was reached. For deep stationary phase timepoints, cells were inoculated and grown for 96 h before collection. When needed, antibiotics were added at the following concentrations: kanamycin 20 μg/mL, chloramphenicol 25 μg/mL, ampicillin 100 μg/mL.

### Hi-C data collection

Cells grown at the desired condition were crosslinked with 10% formaldehyde at room temperature for 30 min, then quenched with 125 mM glycine. Cells were lysed using Ready-Lyse Lysozyme (Epicentre, R1802M) and treated with 0.5% SDS. Solubilized chromatin was digested with Sau3AI for 2 h at 37°C. The digested ends were filled in with Klenow and Biotin-14-dATP, dGTP, dCTP, dTTP. The products were ligated with T4 DNA ligase at 16°C for about 20 h. Crosslinking was reversed at 65°C for about 20 h in the presence of EDTA, proteinase K, and 0.5% SDS. The DNA was then extracted twice with phenol/chloroform/isoamyl alcohol (PCI, 25:24:1), precipitated with ethanol, and resuspended in 20 µl of 0.1×TE buffer. Biotin from non-ligated ends was removed using T4 polymerase (4 h at 20°C), followed by extraction with PCI. The DNA was then sheared by sonication for 12 min with 20% amplitude using a Qsonica Q800R2 water bath sonicator. The sheared DNA was used for library preparation with the NEBNext UltraII kit (E7645). Biotinylated DNA fragments were purified using 10 µL streptavidin beads. DNA-bound beads were used for PCR in a 50 µL reaction for 14 cycles. PCR products were purified using Ampure beads (Beckman, A63881) and sequenced at the Indiana University Center for Genomics and Bioinformatics using NextSeq500 or NextSeq2000. Paired-end sequencing reads were mapped to the genome of *E. coli* W3110 (NCBI reference sequence GCA_000005845.2 with a deletion from 1398953 to 1438196) using the same pipeline described previously^40,54^. The *E. coli* W3110 genome was first divided into 461 10-kb bins. Subsequent analysis and visualization were done using R scripts.

### Hi-C peak finding algorithm

Hi-C heatmaps with a resolution of 10 kb were converted into a 461 × 461, 32-bit unsigned TIFF image and imported into ImageJ^55^. The heat-map was rescaled and converted to a 16-bit signed TIFF file for further analysis (Fig. S1A). We then used a custom ImageJ macro to analyze the strength of the peaks. Because the circular chromosome has no true start or end, the TIFF images were first extended to a size of 661 × 661 pixels, ensuring that the edges of the original image would not be treated differently than the central pixels (Fig. S1B). We next used a 10-pixel (100-kb) radius rolling ball background subtraction to remove features of the Hi-C heat map that varied over larger distance scales (Fig. S1C). After background subtraction (Fig. S1D), we used a threshold to identify the central ridge of self-interactions near the line *y*=*x* (Fig. S1E) and removed these from the image (Fig. S1F). We reduced the shot noise in the resulting image by subtracting a constant set to 2.5 times the mean pixel value. After processing the heatmap, we trimmed the image back to the original resolution of 461×461 pixels (Fig. S1G). The final processed image was integrated along the y-axis to produce a measure of the strength of the peaks over the length of the genome (Fig. S1H). The image was also blurred using a 1-pixel (10 kb) radius Gaussian filter for easier display (Fig. S1I).

### ChIP-seq

Chromatin immunoprecipitation (ChIP) was performed as described previously^40^. Cells were grown in HDA medium at 37°C to the desired growth phase. Cells were crosslinked using 10% formaldehyde for 30 min at room temperature, then quenched using 125 mM glycine, washed with PBS, and lysed using lysozyme. A Qsonica Q800R2 water bath sonicator was used to shear crosslinked chromatin to an average size of 170 bp. The lysate was precleared using Protein A magnetic beads (GE Healthcare/Cytiva 28951378, Marlborough, MA) and then incubated with anti-H-NS^37^ or anti-mCherry^40^ antibodies overnight at 4°C. The following day, the lysate was incubated with Protein A magnetic beads for 1 h at 4°C. After washes and elution, the immunoprecipitate was incubated at 65°C overnight to reverse the crosslinking. The DNA was further treated with RNaseA and Proteinase K, extracted with PCI, resuspended in 100 µL EB, and used for library preparation with the NEBNext Ultra II kit (E7645). Library sequencing was performed using Illumina NextSeq500 or Nextseq2000 (Illumina, San Diego, CA) at the IU Center for Genomics and Bioinformatics. The sequencing reads were mapped to the genome of *E. coli* W3110 (NCBI reference sequence GCA_000005845.2 with a deletion from 1398953 to 1438196) using CLC Genomics Workbench (Qiagen, Hilden, Germany). Sequencing reads were normalized by the total number of reads, plotted, and analyzed using R scripts.

### Whole genome sequencing

The whole genome sequencing experiments were performed as previously described^56^. Cells were grown in HDA medium at 37°C to the desired growth phase. Cells were crosslinked with 10% formaldehyde at room temperature for 30 min, then quenched with 125 mM glycine. Cells were lysed using Ready-Lyse Lysozyme (Epicentre, R1802M) and treated with 1% SDS. Extracted DNA was sonicated using a Qsonica Q800L sonicator for 12 min at 20% amplitude to achieve an average fragment size of 170 bp. The DNA library was prepared using NEBNext Ultra II kit (E7645; NEB) and sequenced using Illumina NextSeq500 or Nextseq2000. Sequencing reads were mapped to the *E. coli* W3110 genome (NCBI reference sequence GCA_000005845.2 with a deletion from 1398953 to 1438196) using CLC Genomics Workbench (Qiagen, Hilden, Germany). The mapped reads were normalized to the total number of reads and used as input for the ChIP samples.

### Comparison of ChIP-seq and Hi-C results

Normalized ChIP-seq reads were binned at 10-kb intervals, identical to the bins used for Hi-C analysis. The binned ChIP-seq reads were then re-normalized to have a mean of zero and a standard deviation of 1. Plots of Hi-C peak strength were also renormalized to have a mean of zero and a standard deviation of 1, facilitating comparison. Scatter plots were generated by plotting different pairs of data against each other, and a linear regression was performed to measure the Pearson correlation coefficient between paired datasets (Igor Pro, WaveMetrics, Lake Oswego, OR, USA).

### Single-molecule tracking, single-particle tracking, and super-resolution imaging

Cells expressing HUα-PAmCherry or H-NS-PAmCherry from the native promoter were grown overnight at 30°C with shaking at 250 rpm in HDA medium and then diluted to OD_600_ = 0.03 – 0.05 in 25 – 50 mL of fresh HDA medium and allowed to incubate at 30°C for 96 h.

Spent medium was prepared by centrifuging 10 mL of saturated culture from each culture flask for 6.5 – 7.5 min at 4,950 – 6,600 × g and syringe filtering the supernatant at least twice using a fresh 0.22-µm syringe filter each time. Spent medium was then used to make 2% (m/v) agarose pads, and 1.5 µL of cells were deposited onto the agarose pad along with 1.5 µL of Fluoresbrite carboxylate YG beads (0.35 µm, Polysciences) that were previously suspended in spent medium (1 – 3 µL as-shipped beads diluted into 1 mL of spent medium). The cells and beads were sandwiched onto the agarose pad by a No. 1 coverslip. For imaging exponential phase cells, fresh HDA medium was used to make agarose pads.

PALM imaging and tracking of H-NS-PAmCherry and HUα-PAmCherry were completed on an Olympus IX-71 microscope with either a 100×1.40 NA or a 100×1.45 NA phase-contrast, oil-immersion objective heated to 30°C with an objective heater (Bioptics). Immersion oil optimized for 30°C (Zeiss) was used to reduce optical aberrations. Photoactivation was performed with a 406-nm laser (Coherent Cube 405-100) with a 0.5-2 W/cm^2^ power density and exposure times optimized for each construct (100-400 ms). PAmCherry fusions were imaged with a 561-nm laser (Coherent Sapphire 561-50) with a power density of 0.46 and 0.55 kW/cm^2^ for HUα-PAmCherry and H-NS-PAmCherry, respectively. Fluorescence emission from PAmCherry was filtered with a 561-nm long-pass filter and imaged with a 512×512-pixel Photometrics Evolve electron-multiplying charge-coupled detector (EMCCD) camera. Each cell was imaged for no longer than 6.5 min to prevent phototoxicity. Each prepared sample of cells on an agarose pad was imaged for no more than 1 h.

eGFP-labelled protein nanocages^44^ were expressed from an arabinose-inducible promoter on plasmid pLAM003^33^. 200 µL of saturated overnight culture grown at 37°C was diluted into 20 mL of fresh HDA medium containing 100 µg/mL spectinomycin and 18-20 µg/mL arabinose, and the culture was incubated for 96 h at 30°C. After the incubation time, spent medium was prepared by centrifuging the culture and filtering the supernatant twice as described above. Then, cells were concentrated 10-fold, and 2 µL of concentrated cells was pipetted directly onto a 2% (w/v) agarose pad made from spent medium and sandwiched by a No. 1 coverslip.

Single-particle nanocage imaging was performed on an Olympus IX-71 inverted microscope with a 100×1.40 NA oil-immersion objective heated to 30°C by an objective heater (Bioptics) and using appropriate index-matched oil. The cell sample was mounted on the microscope objective and allowed to rest for 20 min to thermally equilibrate before imaging. Each sample was imaged for 60 – 75 min. Fluorescence imaging of the nanocage-eGFP fusions was performed with a 488-nm laser (Coherent Sapphire 488-550) with a power density of 160 W/cm^2^. Movies capturing nanocage diffusion were acquired continuously with a 512×512-pixel Photometrics Evolve EMCCD camera (40-ms frame time). Each region was imaged for less than 1 min to reduce phototoxicity.

Single fluorescent molecules or particles within the cell masks were localized in each imaging frame using the SMALL-LABS algorithm^57^. Phase-contrast images of the cells for each growth condition were segmented using the Cellpose package^58^. For each segmented cell, the length and width were calculated from the maximum and minimum Feret diameters, respectively, and the aspect ratio is the ratio of length to width. Cells with 1.5 – 4 µm lengths were selected for further analysis. The coordinates of the fluorescent molecules or particles within each cell were normalized, and 2D histograms were constructed; the heatmaps were symmetrized along the long and short axes and normalized to a total localization probability of one. The aspect ratio used to display each heatmap for each condition indicates the average aspect ratio for each condition (Fig. S7).

### Calculation of the fraction of nanocages excluded from the nucleoid

The nucleoid boundary was determined by the 60% contour line from the HUα-PAmCherry heatmap at each condition, and the apparent excluded fraction, *exc_app_*, is the integral of the normalized protein nanocage occupancy heatmap outside of this nucleoid boundary. However, this 2D projection miscounts the excluded fraction because nanocages above and below the nucleoid are counted as being inside the nucleoid boundary. To adjust for the 3D cell geometry, we transformed the 2D cell and nucleoid contours from the heatmap into 3D based on assuming radial symmetry and rotating about the cell long axis. We then projected a 3D cell with all nanocages in the cytoplasm (no nanocages in the nucleoid) back to 2D to determine the maximum apparent excluded fraction, *exc_max_*, and we projected a 3D cell with equal concentration of nanocages in the cytoplasm and nucleoid back to 2D to determine the corresponding excluded fraction, *exc*_0_. The corrected, normalized excluded fraction, *exc_norm_*, was therefore calculated by comparing the apparent excluded fraction, *exc_app_*, to *excmax* and *exc*_0_:

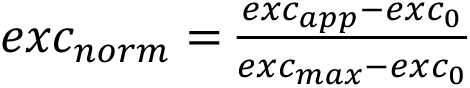

This normalization process yields 0 ≤ *exc_norm_* ≤ 1, such that *exc_norm_* = 1 indicates that no nanocages are within the nucleoid, and *exc_norm_* = 0 indicates equal concentrations of nanocages in the cytoplasm and the nucleoid.

### RNA-seq

RNA was extracted from *E. coli* by adapting a hot-phenol method^59^. Cells were grown in HDA medium at 37°C to the desired growth phase. ∼5 OD units of culture were collected and fixed with an equal volume of methanol. The cells were pelleted, then frozen in liquid nitrogen and stored at -80°C for less than a week before further steps.

Pellets were resuspended in 800 µL hot LETS buffer (0.01 M LiCl, 0.05 M EDTA, 0.01 M Tris pH 7.4, and 1% SDS) heated to 75°C for 1 h. This mixture was then added to 650 mg of acid-washed glass beads (sigma G4649) and 600 µL phenol that had been heated to 75°C and vortexed for 3 min to lyse the cells before 600 µL chloroform was added. The sample was spun at 21,300 × g for 10 min at 4°C. Unless otherwise stated, the same centrifugation speed was used for the following steps. The aqueous layer was collected and added to 800 µL phenol-chloroform (5:1, pH 4.3, heated to 75°C). The sample was vortexed for 3 min and centrifuged for 10 min at 4°C. 700 µL of the aqueous layer was added to 700 µL isopropanol and left at room temperature for 10 min to precipitate the nucleic acids present in the sample. The pellet was collected by spinning at 4°C for 25 minutes, then washed using 1 mL ice-cold ethanol (75%). The sample was spun at 5 min at 4°C to collect the pellet, which was air dried at room temperature for 10 min to remove residual ethanol.

20 µL DEPC-treated water (Thermo Fisher, r0601) was used to resuspend the pellet and the samples were heated to 55°C to resuspend the nucleic acids. 300 µL TRIzol (Thermo Fisher, 1559602) was added to each sample before vortexing for 15 s then incubating at room temperature for 5 min. 60 µL chloroform was added to the tubes, which were mixed by inverting before being placed on ice. After 2 min, the samples were spun at 4°C for 15 min, and 175 µL of the aqueous layer was removed and mixed with 175 µL isopropanol to precipitate the nucleic acids. The samples were incubated at room temperature for 10 min before being centrifuged for 25 min at 4°C. The supernatant was removed, and 250 µL ice-cold 75% ethanol was used to wash the pellet. After spinning for 5 min at 4°C, the supernatant was removed and the pellet air dried at room temperature for 10 min to remove residual ethanol. 20 µL DEPC-treated water was used to resuspend the pellet, which was heated to 55°C to help resuspension.

The TURBO DNA-free kit (Thermo Fisher AM1907) was used to degrade DNA present in the sample. In short, 2 µL 10XTURBO DNase buffer was added to each sample before adding 1 µL TURBO DNase. The tube was incubated at 37°C for 20 min before adding 2 µL DNase Inactivation Reagent to stop the reaction. After incubation at room temperature for 5 min, the sample was spun at 10,000 × g at room temperature for 1.5 min. The supernatant was transferred to a clean tube before performing a final TRIzol extraction as described above to purify the RNA. 40 µL DEPC-treated water was used to resuspend the pellet.

2 µL of the sample was used to run an agarose gel to confirm the presence of RNA. RNA concentration was first estimated using a NanoDrop (Thermo Fisher, ND-One-W) and diluted to ∼200 ng/µL, before loading on an Agilent 2200 Tape Station to better measure the RNA concentration and integrity. The RNA concentrations of our sample were between 216 ng/µl and 477 ng/µl, and the RNA integrity number (RIN) values were between 8.8 and 10.0. The diluted RNA was sent to SeqCenter, LLC in Pittsburgh, PA for rRNA depletion, library preparation, and paired-end sequencing on the Illumina NovaSeq X Plus platform. Each sample had 6.8 to 13 million paired-end reads. Reads were mapped to the *E. coli* W3110 genome (NCBI reference sequence GCA_000005845.2 with a deletion from 1398953 to 1438196) using CLC Genomics Workbench (Qiagen, Hilden, Germany).

### RNA-seq analysis

RNA-seq reads were imported into the Galaxy server^60^ and processed using the EdgeR package^42^. To visualize the spatial distribution of upregulated and downregulated genes in paired datasets, we selected all genes with a p-value < 0.05 in both datasets and plotted the log_2_ of the fold change (FC) against the transcription start site of each gene (Igor Pro, WaveMetrics, Lake Oswego, OR, USA). To compare RNA-seq data with other datasets, we generated a weighted histogram of downregulated genes as follows. First, the genome was divided into 461 10-kb bins. Then, for each gene with FC< 0.5, we added a value of -log_2_(FC) into that bin. The final weighted histogram was then renormalized to have a mean of zero and a standard deviation of 1 (Igor Pro, WaveMetrics, Lake Oswego, OR, USA).

### Plasmid construction

**pWX1209** [IPTG-dependent p15a *ori* to express *flp, amp*] was built through the isothermal assembly of two fragments: 1) pSW439^61^ was digested with BamHI and HindIII and gel purified to render the plasmid backbone; 2) pCP20^62^ was amplified with oWX3405 and oWX3406 to give *flp* recombinase. The plasmid was verified by whole plasmid sequencing.

### Strain construction

**cWX2356** [*TB10, Δfis FRT-kan-FRT*] was generated through lambda recombineering. The *FRT-kan-FRT* cassette was amplified from pKD4^63^ using primers oWX2811 and oWX2812 with homology upstream and downstream of *fis*. The resulting PCR product was treated using DpnI, column-purified, and electroporated into TB10^64^. The construct was amplified and sequenced using oWX2777 and oWX2780.

**cWX2471** [*W3110, Δfis FRT-kan-FRT*] was generated by P1 transduction of W3110 with P1 lysate from strain cWX2356.

**cWX2478** [*W3110, Δfis FRT-kan-FRT*] was generated by P1 transduction of W3110 with P1 lysate from strain cWX2471. This second step of transduction was to ensure that Δ*fis* was the only mutation in this strain.

**cWX2870** [W3110, *Δhns FRT-kan-FRT*] was generated by P1 transduction of W3110 with P1 lysate from JW1225-KC in the Keio collection^65^.

**cWX2876** [W3110, *ΔihfA FRT-kan-FRT*] was generated by P1 transduction of W3110 with P1 lysate from JW1702-KC in the Keio collection^65^.

**cWX2878** [W3110, *ΔihfB FRT-kan-FRT*] was generated by P1 transduction of W3110 with P1 lysate JW0895-KC in the Keio collection^65^.

**cWX2882** [*W3110, hns-PAmCherry FRT-cam-FRT*] was generated by P1 transduction of W3110 with P1 lysate from strain *hns-PAmCherry-cam*^39^.

**cWX2886** [*W3110, Δhns FRT*] was generated by FLP-mediated site-specific excision of the *FRT-kan-FRT* from cWX2870 using pCP20^62^. An *FRT* scar was left after excision of the cassette.

**cWX2889** [*W3110, Δhns FRT, hupA-PAmCherry FRT-cam-FRT*] was generated by P1 transduction of cWX2886 with P1 lysate from *hupA-PAmCherry FRT-cam-FRT*^39^.

**cWX2934** [*W3110, ΔihfA FRT*] was generated by FLP-mediated site-specific excision of the *FRT-kan-FRT* from cWX2876 using pWX1209. The transformants were plated on LB agar containing 100 mg/mL of ampicillin and 500 mM IPTG. Transformants were streaked on LB agar to cure pWX1209. Loss of the cassette and pWX1209 was verified by sensitivity to kanamycin and ampicillin. pWX1209 was used for excision rather than pCP20^62^ because *ihf* is required for replication of pCP20, which has pSC101 origin. An *FRT* scar was left after excision of the cassette.

**cWX2943** [*W3110, ΔihfA FRT, ΔihfB FRT-kan-FRT*] was generated by P1 transduction of cWX2934 with P1 lysate from strain cWX2878.

**cWX3338** [*W3110, ΔstpA FRT-kan-FRT*] was generated by P1 transduction of W3110 with P1 lysate from JW2644-KC in the Keio collection^65^.

## Acknowledgements

We thank Christine Jacobs-Wagner for the nanocage plasmids, Xiaowei Zhuang for the HUα-PAmCherry and H-NS-PAmCherry strains, Robert Landick for H-NS antibodies, David Rudner for mCherry antibodies, Daniel Foust for assistance with data analysis, Xheni Karaboja for technical assistance, Dan Kearns, Tom Bernhardt, and Rodrigo Reyes-Lamothe for protocols, Cristina Landeta and James McKinlay for strains, Indiana University Center for Genomics and Bioinformatics for high-throughput DNA sequencing, and SeqCenter for RNA-seq.

## Author Contributions

L.E.W. constructed plasmids and strains, performed ChIP-seq, Hi-C, RNA-seq experiments and analysis. X.D. performed nanocage single-particle tracking experiments and data analysis. E.E.W. developed RNA-seq procedure and analysis. L.A.M. performed H-NS single-molecule tracking experiments and analysis. Z.R. and G.G.H. constructed strains, performed ChIP-seq and Hi-C experiments. D.E.H.F. performed HUα super-resolution microscopy. L.H. constructed plasmids and strains. E.A. analyzed ChIP-seq, Hi-C and RNA-seq data. A.S.M, J.S.B. and X.W. supervised the study. L.E.W, X.D., E.E.W., L.A.M, E.A., J.S.B. and X.W. wrote the text with input from all authors.

## Materials Availability

Plasmids and strains generated in this study are available from the corresponding authors with a completed Materials Transfer Agreement.

## Data Availability

Hi-C, ChIP-seq, and WGS data were deposited to the NCBI Gene Expression Omnibus.

## Code Availability

Data analysis tools are available on the Biteen Lab GitHub at https://github.com/BiteenMatlab. Any additional information required to analyze the data reported in this paper is available from the corresponding authors upon request.

---

**Figure S1.**
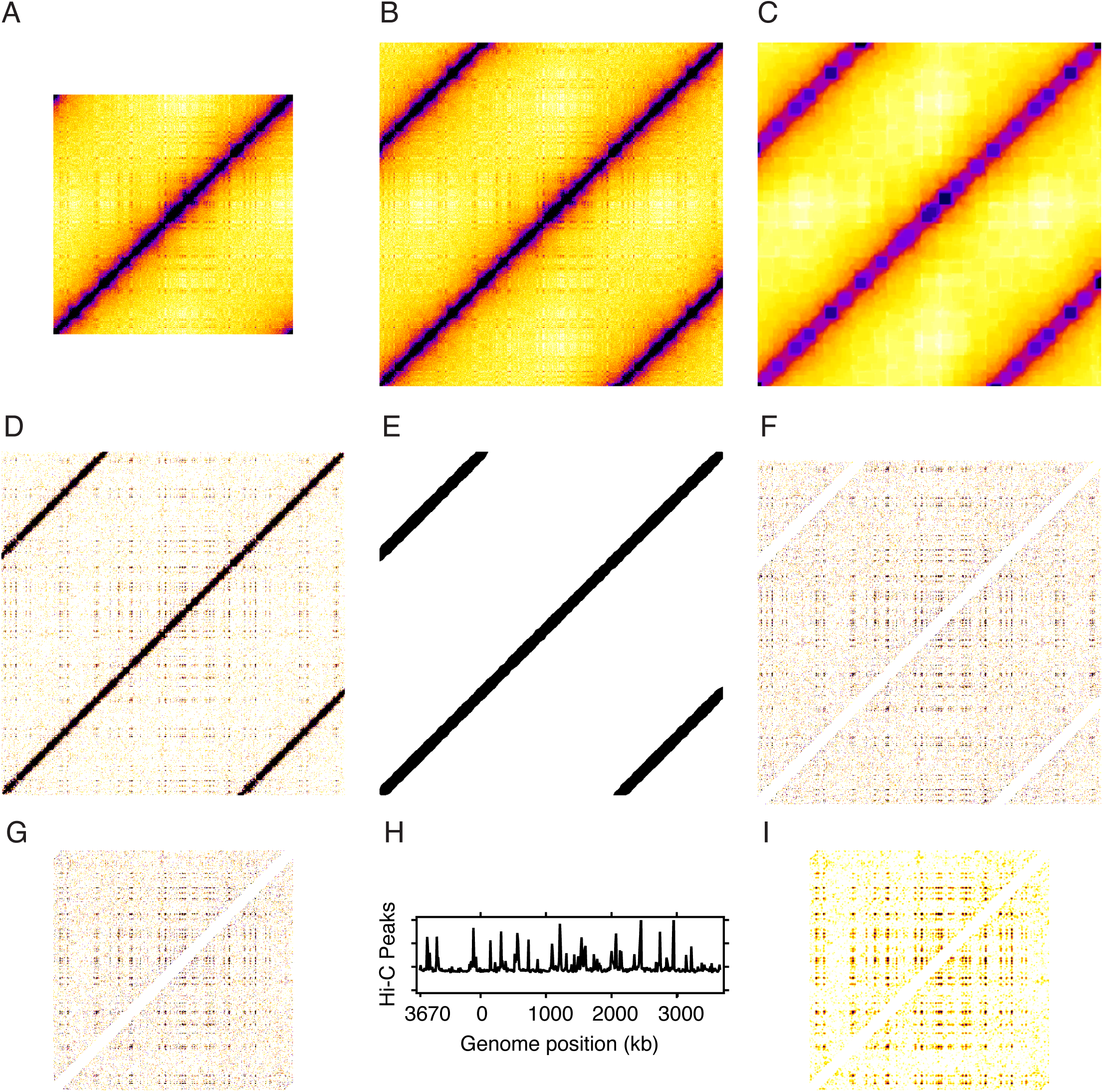
Peak finding algorithm used to identify long-range interactions. **(A)** The Hi-C density plot at 10-kb resolution is converted to an unsigned 32-bit TIFF file and loaded into ImageJ. **(B)** The image is extended with repeated information to remove edge effects during subsequent analysis. **(C)** A 10-pixel (100-kb) radius rolling ball is applied to the image to estimate the slowly varying background components of the Hi-C plot. **(D)** This background is subtracted from the image in (B) to isolate the sharp peaks. **(E)** The image from (D) is blurred, and an automated threshold is applied to identify the ridge corresponding to self-interaction. **(F)** The thresholded image from (E) is used to remove the ridge from (D), producing an image with only sharp peaks remaining. **(G)** The image from (F) is trimmed back to its original size. **(H)** The image from (G) is integrated to determine the linear Hi-C peak density. **(I)** A 1-pixel (10-kb) Gaussian blur is applied to the image from (G) to make it easier to identify the peaks when printed at a compact size.

**Figure S2.**
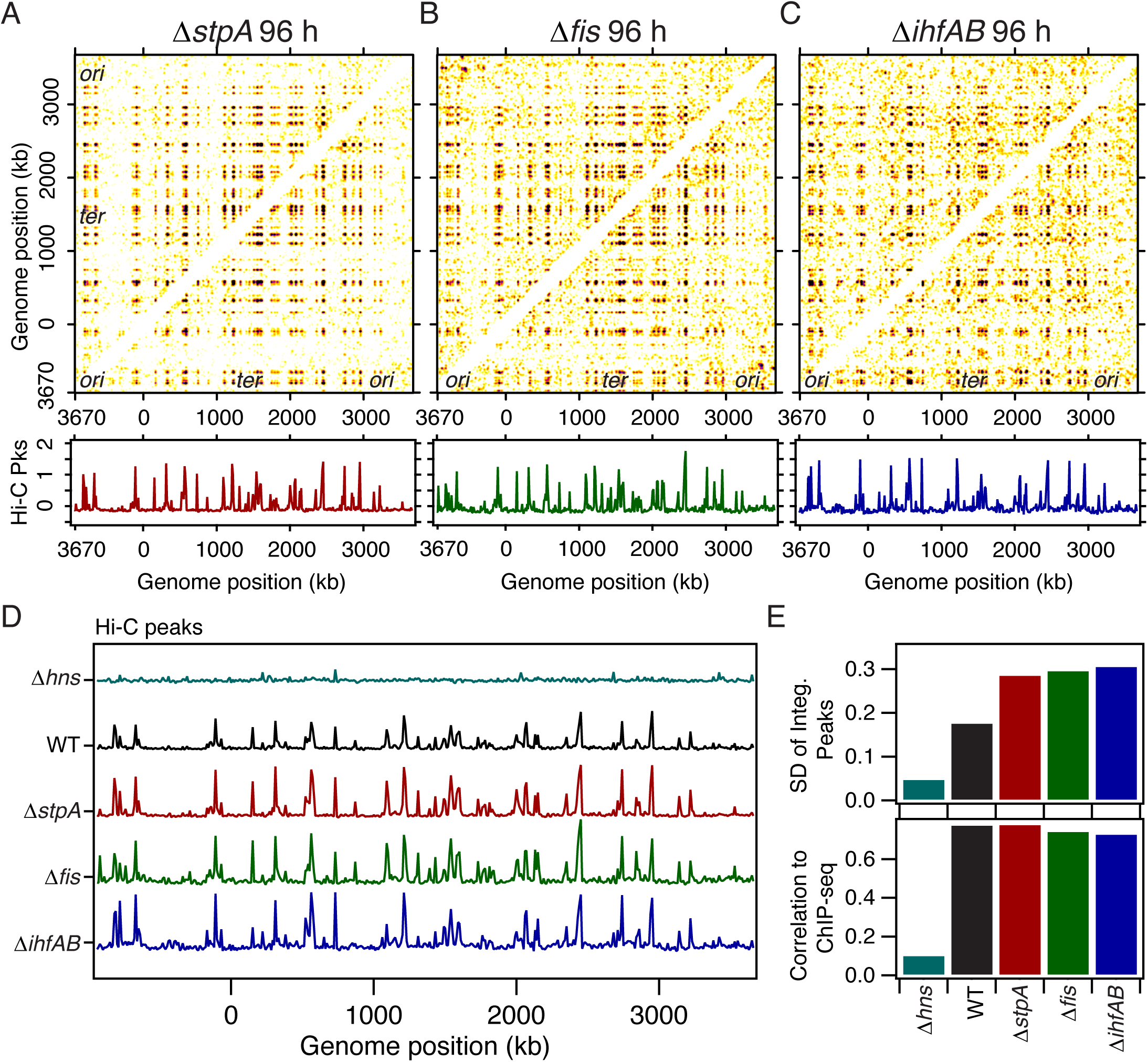
Peak finding analysis for indicated mutants at 96h. **(A-C)** Analysis of Hi-C loop anchors using the rolling ball background subtraction algorithm for *ΔstpA* (cWX3338, A), *Δfis* (cWX2478, B), and *ΔihfAB* (cWX2943, C) during deep stationary phase. Top panels show the genome-wide Hi-C peaks. Bottom panels show the linear integrated density of the peaks across the genome. See Method Details and Fig. S1. **(D)** Comparison of the integrated densities of Hi-C peaks from different strains during deep stationary phase. **(E)** Comparison of the standard deviation (SD) of the integrated Hi-C peak densities (upper) and correlation to the anti-H-NS ChIP-seq peaks at 96 h (lower), demonstrating the relative strength of the peaks in each condition and their relation to H-NS binding sites.

**Figure S3.**
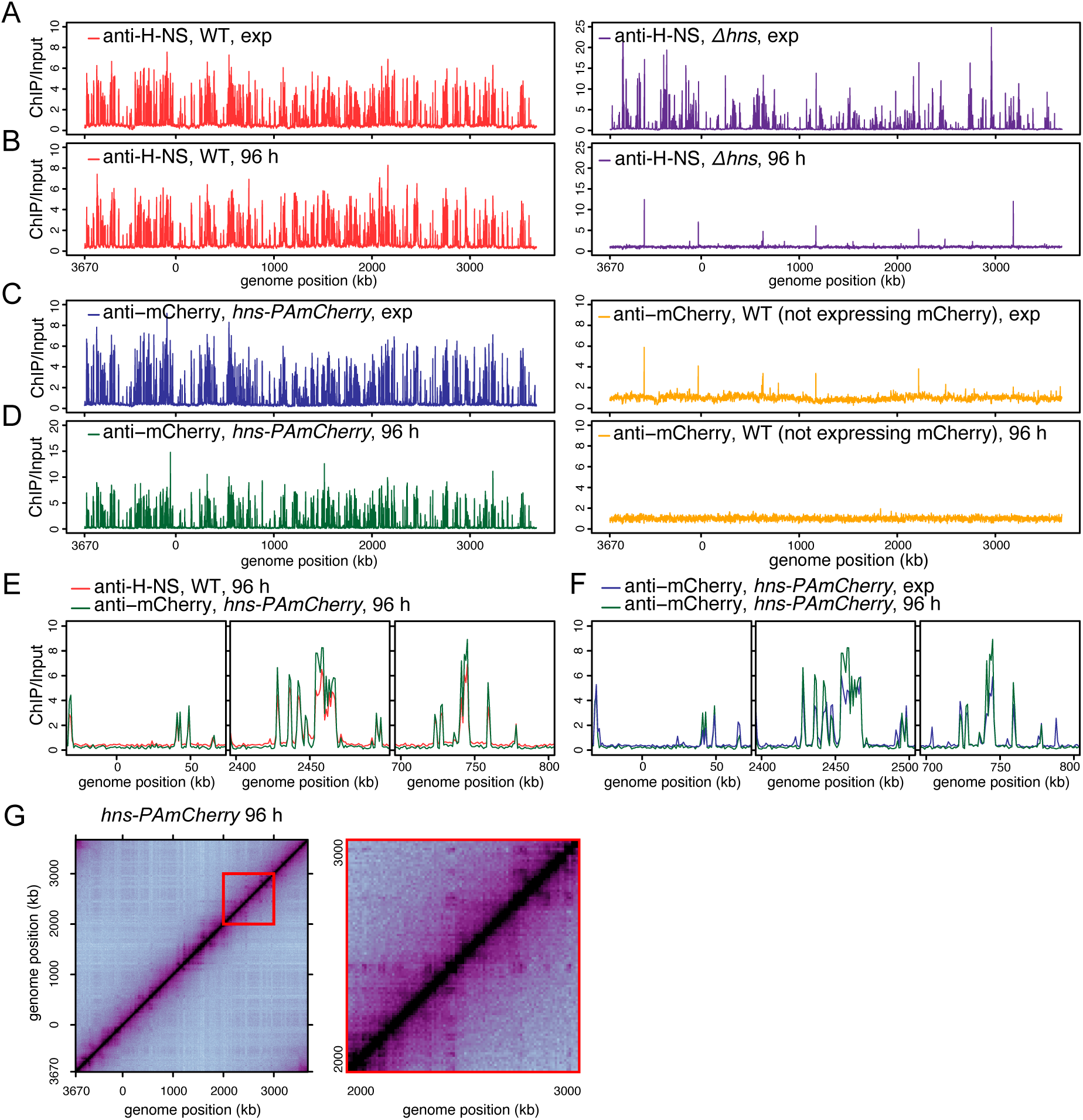
ChIP-seq results using anti-H-NS and anti-mCherry antibodies. **(A-B)** Anti-H-NS ChIP-seq in the WT strain (left) or *Δhns* (cWX2870, right) at exponential phase (A) or deep stationary phase (B). The sequencing reads at each base pair were normalized to the total number of reads. Then, ChIP enrichments were plotted by calculating ChIP/input. The genome was binned at 1-kb resolution and oriented to start at *oriC*. **(C-D)** Anti-mCherry ChIP-seq in *hns-PAmCherry* (cWX2882 left) or the WT (i.e., not expressing mCherry, right) at exponential phase (C) or deep stationary phase (D). **(E)** Overlay of anti-H-NS ChIP-seq in the WT and anti-mCherry ChIP-seq in *hns-PAmCherry* (cWX2882). Cells were grown during deep stationary phase. Three representative regions are shown: the left panel is not a loop anchor identified by Hi-C, while the middle and right panels are regions identified as loop anchors by Hi-C. **(F)** Anti-mCherry ChIP-seq in *hns-PAmCherry* (cWX2882) grown at exponential phase and deep stationary phase for the three representative regions shown in (E). **(G)** Normalized Hi-C results for *hns-PAmCherry* (cWX2882) grown at deep stationary phase. Left, whole-genome Hi-C map. Right, zoom-in to a 1000-kb region (red box).

**Figure S4.**
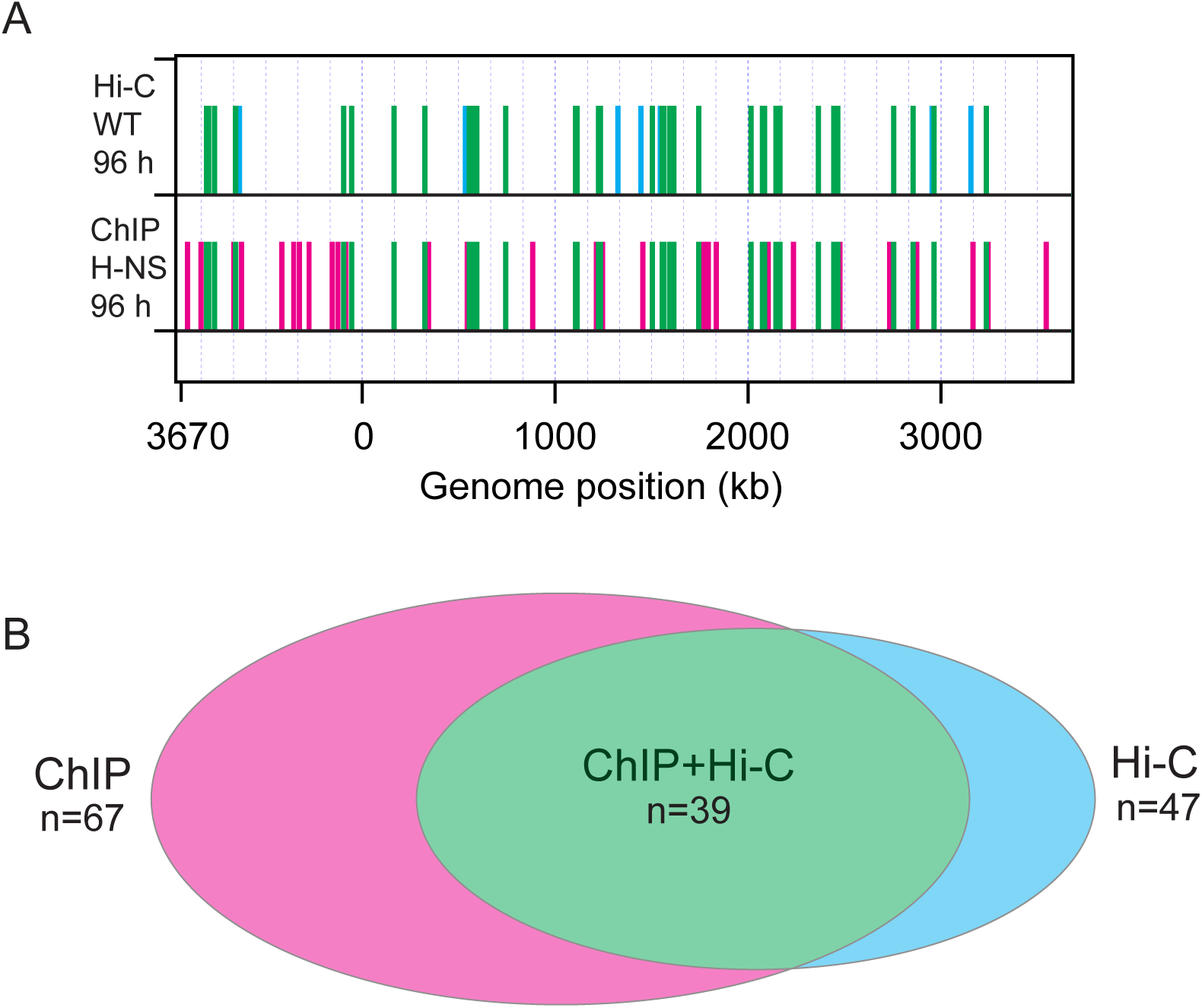
Comparison of ChIP-seq and Hi-C datasets for 96-h samples. **(A)** Binary representation of the 96 h Hi-C peaks and ChIP-seq densities plotted in Fig. 2D (2^nd^ and 4^th^ panels). Any bin greater than one standard deviation above the mean is assigned a value of one; all other bins are assigned a value of 0. Bins that are assigned a value of 1 in the Hi-C and ChIP-seq datasets are colored green, bins only assigned a value of 1 in the Hi-C dataset are colored cyan, and bins only assigned a value of 1 in the ChIP-seq dataset are colored magenta. **(B)** Venn diagram showing the number of bins that overlap between the two datasets plotted in (A).

**Figure S5.**
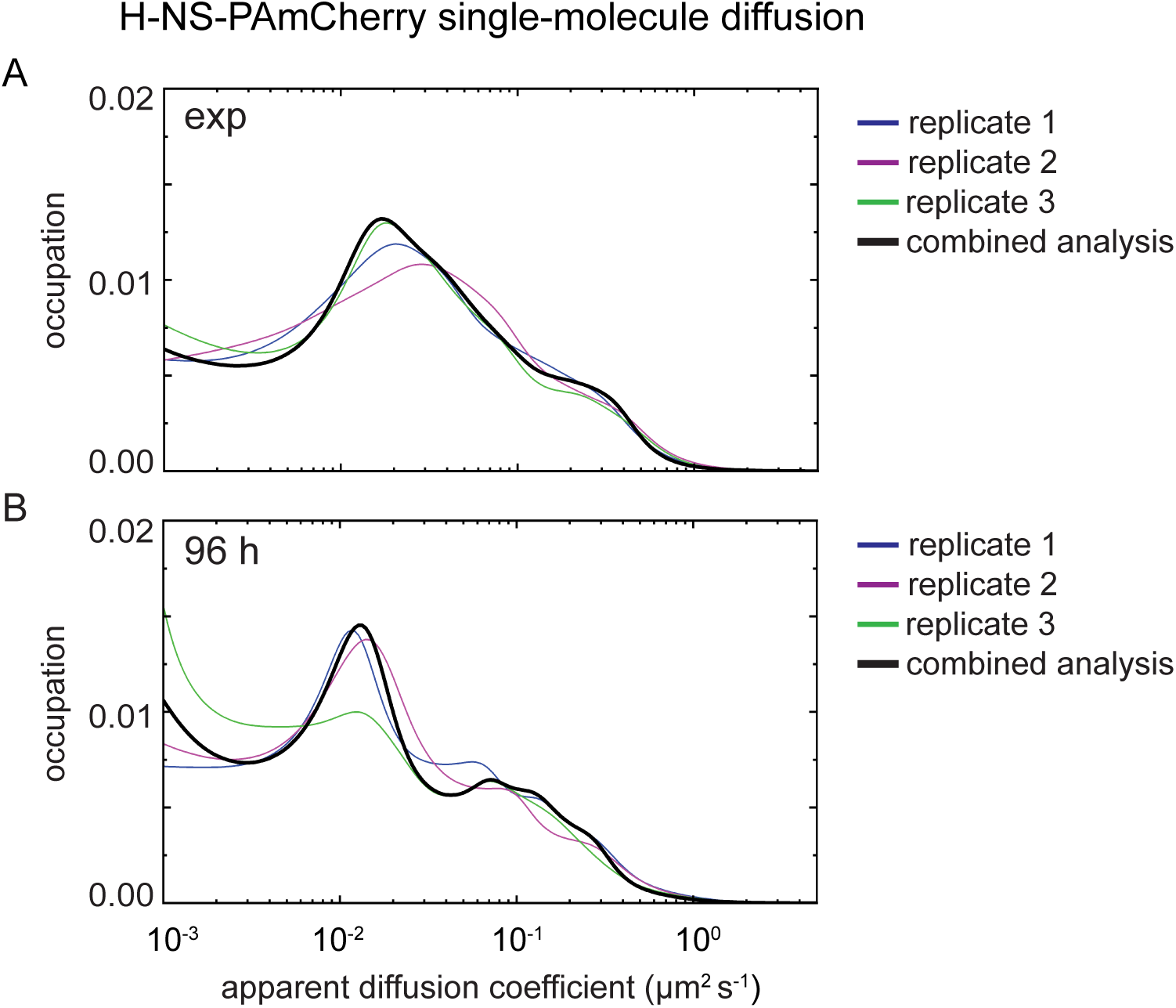
H-NS-PAmCherry dynamics are reproducible across biological replicates. Apparent diffusion coefficient distributions of H-NS-PAmCherry (cWX2882) measured in the exponential phase (A) or deep stationary phase (B) for three biological replicates. Each biological replicate contains single-molecule tracking data for *N* ≥ 7 cells with *n* ∼ 700 trajectories per replicate. The black line is the result of applying the State Array Bayesian framework^66^ to data pooled from all three biological replicates. Variance in the diffusion analysis among biological replicates is small relative to the difference in H-NS-PAmCherry dynamics measured between the exponential and stationary phases.

**Figure S6.**
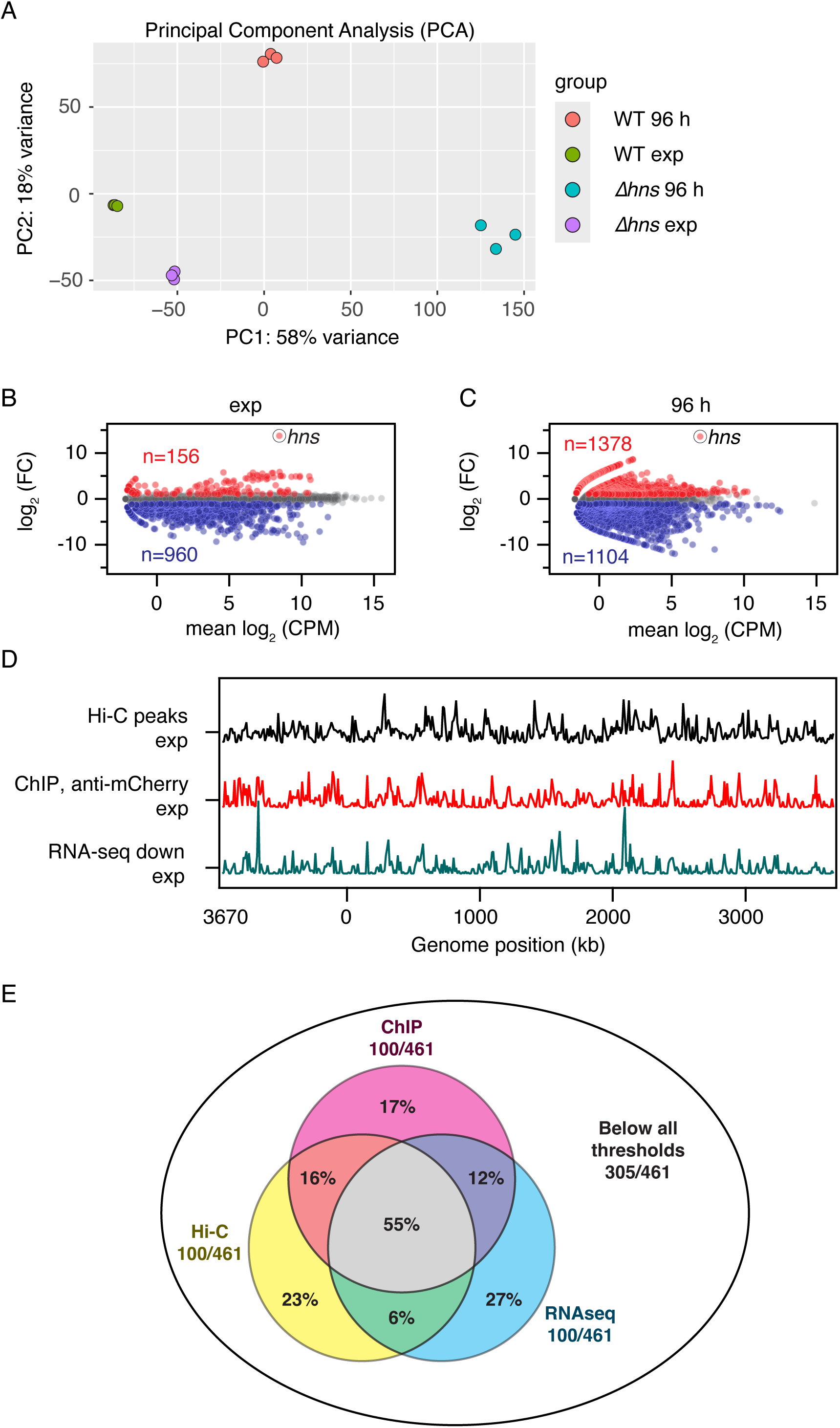
RNA-seq analysis. (A) Principal component analysis (PCA) of the three biological replicates of RNA-seq data for WT and *Δhns* (cWX2882) grown at exponential phase or deep stationary phase. **(B-C)** Differential expression of genes in WT and *Δhns* (cWX2882) at exponential phase (B) and deep stationary phase (C). For each gene, the mean expression in the WT strain was divided by the corresponding value in the *Δhns* strain to calculate fold change (FC=WT*/Δhns*). The y-axis represents log_2_ of FC. The x-axis represents log_2_ of the counts per million bases (CPM) for each gene. Gray dots represent |log_2_(FC)|< 1. Blue and red dots represent log_2_(FC) <-1 and or >1, respectively. The *hns* gene is highlighted. Number (*n*) of downregulated (blue) or upregulated (red) genes are shown. **(D)** Comparison of Hi-C, ChIP-seq, and RNA-seq results in 10-kb bins in exponential phase. Correlations between the different sets are relatively weak in exponential phase (Hi-C vs. ChIP-seq: ρ = 0.11; Hi-C vs. RNA-seq: ρ = 0.20; RNA-seq vs. ChIP-seq: ρ = 0.58). **(E)** Venn diagram showing the spatial overlap between the 100 highest bins from the 96-h datasets in Fig. 4D. More than half of all high bins in each dataset overlap with high bins in both of the other datasets.

**Figure S7.**
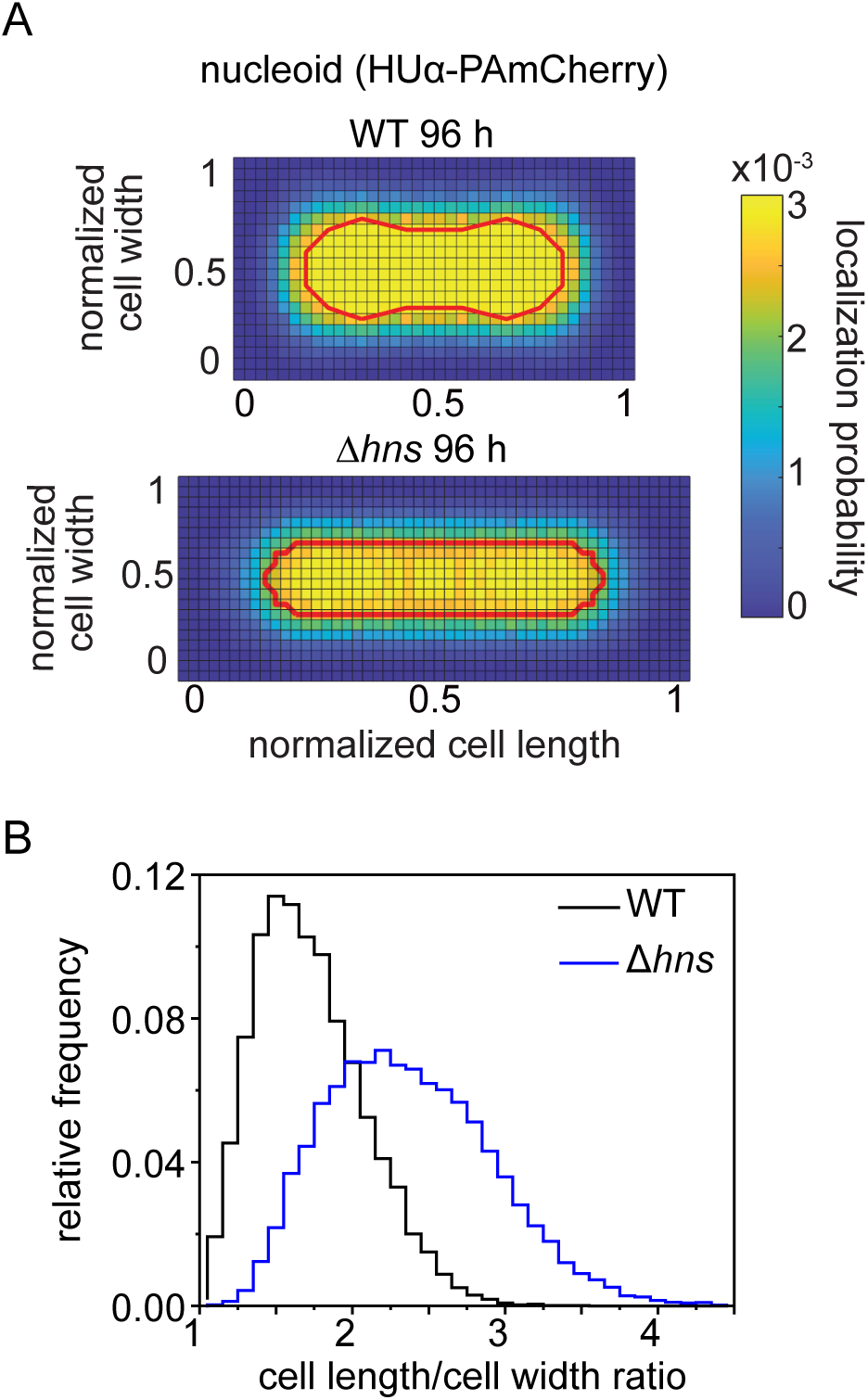
HUα-PAmCherry super-resolution heatmap and cell aspect ratio distribution. **(A)** HUα-PAmCherry localization heatmap of WT and Δ*hns* cells at deep stationary phase (96 h). Red line represents top 60% contour line of localization probability, defining the nucleoid region. **(B)** Cell aspect ratio of WT and Δ*hns* cells at deep stationary phase. The aspect ratio of each cell is the ratio of the maximum and minimum Feret diameters of each cell mask. The dataset comprises >10000 cells segmented by the Cellpose algorithm^58^.

